# Metabolic deregulation in prostate cancer

**DOI:** 10.1101/371567

**Authors:** Sriganesh Srihari, Paula Tattam, Rebecca Simpson, Elliot Smith

## Abstract

**Introduction:** The prostate exhibits a unique metabolism that changes during initial neoplasia to aggressive prostate cancer (PCa) and metastasis. The study of PCa metabolism thus represents a new avenue for diagnostics, particularly early diagnosis of aggressive PCa cases.

**Results:** Here, using transcriptomics data from The Cancer Genome Atlas (498 PCa patients), we identified six metabolic subgroups (C1-C6) of PCa that showed distinct disease-free survival outcomes (p<0.0001). In particular, we identified at least two PCa subgroups (C5 and C3) that exhibited significant poor prognosis (~70% and 30-40% relapse by the first 72 months; hazards ratios 9.4 and 4.4, respectively, relative to the best prognosis cluster C4 that showed <20% relapse even by 120 months). The subgroups were reproducible in an independent dataset from Taylors et al. 2010 (215 patients; p=0.00088). The subgroups displayed distinct metabolic profiles vis-à-vis normal tissues; measured as ‘deregulation’ of metabolic pathways (using Pathifier, Drier & Domany, 2013). In particular, the poor-prognosis subgroups C5 and C3 showed considerable deregulation for pathways involved in synthesis and catabolism of complex forms of lipids and carbohydrates, amino acids, and TCA cycle, and these were exhibited in parallel or in the face of glycolysis, a common form of energy production in cancer cells. Furthermore, the subgroups were significantly over-enriched for different sets of genetic alterations [particularly, deletions/mutations in BRCA1 and TP53 (C5), RB1 and STK11(C3); and AR amplifications (C1); p≤8.6E-04], suggesting that distinct alterations may be underpinning the subgroups and ‘pushing’ the subgroups towards their unique metabolic profiles. Finally, applying the classifier to blood expression profiles from 42 active surveillance (AS) and 65 advanced castrate resistant PCa (ACRPC) patients determined based on prostate-specific antigen (PSA) levels (Olmos *et al.*, 2012) assigned 70.77% ACPRC, and interestingly reassigned 59.52% AS patients to at least one of the poor prognosis subgroups (C5, C3) with 35.71% to the poor and metabolically deregulated subgroup C3.

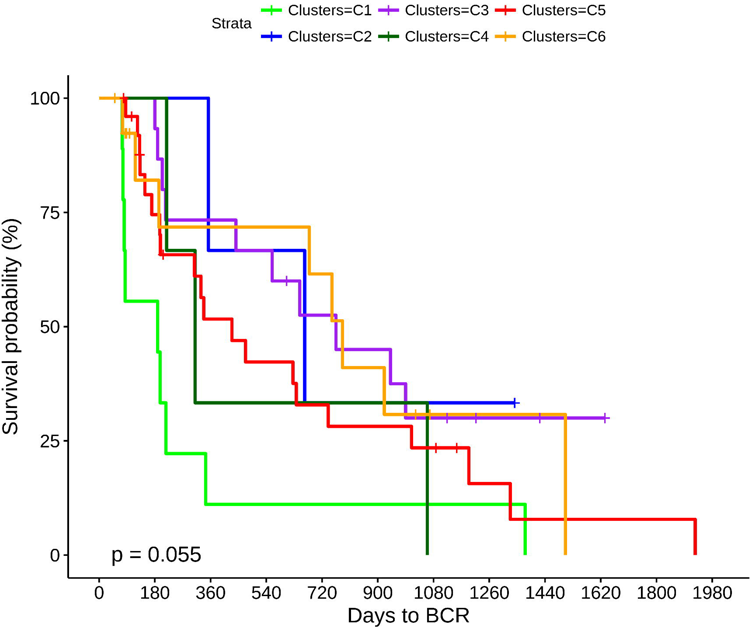

**Conclusion:** The identification of PCa subgroups displaying distinct clinical outcomes solely from metabolic expression profiles of PCa tumours reiterates the significant link between deregulated metabolism and PCa outcomes (Eidelman *et al.*, 2017). On the other hand, the time to biochemical relapse (rise in PSA levels) was not indicative of the early relapse seen for the metabolically deregulated subgroups C3 and C5 (these show considerably late BCR compared to C4). Our study thus highlights specific processes (elevated lipid and carbohydrate metabolism pathways) that could be better indicators than PSA for early diagnosis of aggressive PCa.

## Introduction

**Prostate cancer (PCa)** is the second most common type of cancer among males in Western countries [1], with an estimated ~165,000 new cases and ~29,500 cancer deaths in 2018 alone [2]. Fortunately, 80% of patients with an early diagnosis have a good prognosis and radical prostatectomy is still the most adopted approach with a high rate of success. However, prognosis becomes worse if the disease develops late or is diagnosed late and becomes metastatic.

A widely used prostate marker is the prostate-specific antigen (PSA) measured from the blood which is shown to be elevated in many men with prostatic disease [3]. PSA testing is still used as an early screening method to determine likelihood and aggressive phenotype of prostate cancer. Recently, this method has undergone some scrutiny as it may lead to overtreatment in patients who may have been treated sufficiently with active surveillance [4,5]. Other studies have suggested that PSA may be a poor indicator of, and perhaps inversely related to, aggressive disease in obese men [6]. Improved tools for better risk stratification of prostate cancer patients are thus needed.

The prostate exhibits a unique **metabolism** that changes during initial neoplasia to aggressive PCa and metastasis [4]. The study of metabolism of PCa thus represents a new avenue for diagnostics, particularly early diagnosis of aggressive PCa cases [7]. By better understanding the metabolism of prostate cancer, it may be possible to elucidate metabolic biomarkers whose levels may help to diagnose aggressive prostate cancers.

Here, we sought to understand the metabolic profiles of PCa tumours that show different relapse outcomes. Due to lack of well-defined metabolic PCa subtypes (except perhaps those driven by androgen receptor signalling), we first stratified ~500 PCa patients from The Cancer Genome Atlas (TCGA) [8] into **six metabolic subgroups** that were clinically distinct (C1-C6, that display distinct disease-free survival outcomes) and reproducible in an independent dataset from Taylors et al. (215 patients) [9]. We next computed ‘**deregulation’ for 20 metabolic pathways** for the six subgroups using the Pathifier tool [11]. The poor prognosis subgroups displayed considerably evelated synthesis and catabolism of complex forms of lipids (sphingolipids and glycosphingolipids) and carbohydrates (glycosaminoglycans) in parallel to or in the face of glucose and galactose metabolism that are thought to be commons form of energy production in cancer cells [22, 23]. Furthermore, we found that the PCa metabolic subtypes were enriched for genetic alterations in distinct sets of genes (including ATM, BRCA1, PTEN, AR, STK11, and ERG-TMPRSS2), suggesting that distinct alterations may be underpinning the different subtypes and ‘pushing’ the subtypes towards their unique metabolic profiles.

The identification of PCa subgroups displaying different disease-free survival outcomes solely by clustering metabolic expression profiles of prostate tumours reiterates the significant link between deregulated metabolism and PCa outcomes [4,7]. In particular, we find the poor-prognosis PCa subgroups are characterized by markedly deregulated lipid and carbohydrate pathways. On the other hand, the the time to biochemical relapse (BCR, rise in PSA levels) is not indicative of the poor prognosis (C3 and C5 show considerably late BCR compared to the best prognosis subgroups C1 and C4). Therefore, our study highlights specific processes (elevated lipid and carbohydrate metabolism pathways) that could be better indicators than PSA for early diagnosis of aggressive PCa.

## Datasets and Methods

### Molecular and clinical data of PCa patients

RNAseq expression, DNA mutation and copy-number data of genes along with clinical profiles of patients were downloaded for 499 PCa primary solid tumour samples and 52 normal tissue samples from TCGA (498 patients) [8] via the Broad GDAC Firehose (https://gdac.broadinstitute.org/) [12]. RNAseq data was used for clustering patients and the patient clusters were subsequently assessed for differences in disease-free survival and for enrichment of known genetic alterations (mutations and copy number changes of genes). The clinical profiles include data on age at diagnosis, PSA levels, time to disease relapse, and time to biochemical relapse (BCR). Note here that the normal tissue samples are also biopsied from PCa patients. Independent validation was performed by predicting clusters on a dataset from Taylors et al. (2010) which contained RNAseq, DNA copy number and clinical profiles from 216 PCa samples (215 patients).

### List of genes in pathways

In all, 20 pathways were assessed (**Table S1)**. These pathways were assembled from the curated lists by Peng et al. (2018) [13], Fabregat et al. (2016) [14], our previous comprehensive curation of pathways [17], and from KEGG [15] via the InnateDB platform (http://www.innatedb.com/) [16].

#### Metabolism pathways I

Clustering of PCa patients and initial analysis of pathways was performed using genes from amino acid metabolism (348), carbohydrate metabolism (286), lipid metabolism (766), tricarboxylic acid cycle (TCA cycle, 148), and vitamin & cofactor metabolism (168) pathways, the five main pathways listed in Peng et al.[13].

#### Metabolism pathways II

Subsequent breakdown of the analysis from above pathways were performed using:

(I) <u>Carbohydrate metabolism</u>: glycolysis (67), galactose metabolism (31), pentose phosphate metabolism (28), glycosaminoglycan synthesis (21), and glycosaminoglycan degradation (19); and
(ii) <u>Lipid metabolism:</u> sphingolipid metabolism (47), fatty acid elongation (23), fatty acid degradation (44), and unsaturated fatty acid biosynthesis (21).

#### Metabolic pathways III

Mitochondrial subunits of ribosome (133), oxidative phosphorylation (OXPHOS, 137), AMPK/LKB1 (14), and HIF1-a (16).

#### DNA-damage response (DDR) pathways

Homologous recombination (HR, 82), non-homologous end-joining (NHEJ, 15), and mismatch repair (21).

### Identification of patient clusters from metabolic profiles

Z-score normalized RNASeq expression levels for genes in metabolic pathways I (above) were used to identify patient subgroups. Only genes that were over- or under-expressed (>=2 or <=-2 for z-score values; similar to the normalization and thresholding used by Cbioportal [18]) in at least 10% samples (giving 124 genes) were used for clustering the samples. Hierarchical clustering (using hclust and pheatmap package in R) to generate sample subgroups/clusters of sizes *k*=2 to 10 were tried, and the clustering that gave the most significant separation (logrank test p-value) in terms of disease-free survival for the clusters was retained; we call these clusters **C** = {C_1_,…,C_k_}.

### Plotting disease-free/recurrent-free survival curves

Survival curves for recurrence/relapse of disease (time as months) were plotted using the survminer package in R using recurrence/relapse as the endpoint and by censoring out patients who were disease-free. Hazard ratios for the association between disease-free survival and the clusters were computed based on a Cox proportional hazards regression model using the survminer package in R.

### Prediction model for patient clusters

We trained a multinomial (multi-class) logistic regression classifier in R to learn the cluster labels from the TCGA dataset and applied it to predict cluster labels for patients from an independent dataset from Taylors et al., 2010. The features (genes with RNAseq expression values) for the classifier were selected as follows.

For each cluster *C*_i_ in **C** = {*C*_1_,…,*C*_k_}, we first trained a binomial logistic regression classifier on the TCGA dataset to classify *C*_i_ from the remaining clusters **C** – {*C*_i_} and validated the classifier using five-fold (80%-20%) cross-validation. The classifiers had to achieve a minimum AUC of 0.70. We used the genes *F*_i_ that were identified by the classifier as sigificantly associated (p≤0.01) with the probability of the patients to be in class *C*_i_. Our final list of genes (features) **F** was the union **F** = U^1^_k_ *F*_i_. A multinomial classifier was then trained on 80% TCGA data using features **F** and was validated on the remaining 20% using five-fold cross-validation. Subsequently, the classifier was trained on 100% TCGA data and applied on the Taylors et al. dataset. Disease-free survival curves were computed for the predicted clusters to assess the reproducibility of clinical differences between the clusters.

### Assessing deregulation of pathways in individual patients using Pathifier [**11**]

‘Deregulation’ or ‘activity’ of pathways in individual tumour samples relative to normal tissue samples was computed using the Pathifier tool, available as a R package (https://bioconductor.org/packages/release/bioc/html/pathifier.html) [11]. Given the expression levels of a group of genes (e.g. in a pathway), Pathifier computes a ‘deregulation score’ for the pathway in each tumour sample relative to a collection of normal samples. Pathifier computes this score by determining the principal components (PCs) along which the expression levels of the pathway genes vary the most and then plots each sample in this PC space (resulting in a ‘sample cloud’ in the PC space). Pathifier then fits a principal curve [19] that optimally passes through this sample cloud and so the normals (being the baseline) tend to appear at the beginning of the curve whereas the most deregulated samples appear farther along the curve. The deregulation score of the pathway for a sample is then determined by the distance of the sample from the normals along this curve. Pathifier thus provides a <u>single score for each pathway in</u> <u>each sample</u>, which we then use to measure the extent of deregulation (increased or decreased ‘activity’) of the 20 pathways in tumour clusters relative to normals.

### Enrichment for genetic alterations in patient clusters

We estimate whether a patient cluster *C* is significantly enriched for alterations observed for a gene *g* (mutation, amplification and deletion at DNA copy number level, and over- and under-expression at mRNA and protein levels) using a hypergeometric test, as follows:

If *X* is a random variable that follows the hypergeometric distribution and measures the number of successes (alterations in gene *g*) in cluster *C*, then the enrichment p-value for *g*’s alterations in *C* is:

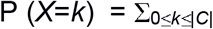

where
*k* = number of alterations for *g* in *C*,
*n* = number of alterations for *g* in the entire population, and
*N* = the population size (498 patients).

We consider P≤0.05 as statistically significant enrichment for *g*’s alterations in *C.*

## Results

### Six ‘metabolic clusters’ of PCa with distinct disease-free survival outcomes

Hierarchical clustering of 498 TCGA PCa patients based on expression profiles of genes from metabolic pathways-I that are altered (over- or under-expressed) in at least 10% patients (124 genes; **Figure S1 & Table S2**) resulted in six patient clusters: C1-C6 (C1: 115 samples, C2: 49, C3: 125, C4: 26, C5: 107, C6: 75) (**Tables 1 and S2**). The six clusters showed significantly different disease-free survival outcomes (logrank test p<0.0001) with clusters C5, C6, C3 (in that order) displaying the quickest time to relapse (**Figure 1a**). In particular, we identified a considerably poor-prognosis cluster C5 that showed recurrence for ~70% patients by 72 months (6 years). 30-40% patients in C6 and C3 showed relapse by ~72 months. Cluster C4 showed the slowest time to relapse –<20% relapse even by 120 months. Cox-proportional hazard ratios were significantly high for clusters C5, C6, and C3, relative to normal – C5 (HR=22, p=0.002), C6 (HR=14.4, p=0.009), and C3 (HR=10, p=0.024).

**Figure 1.**
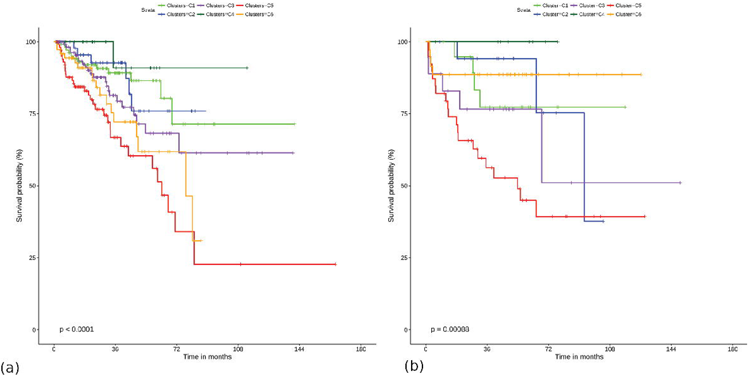
Six clinically distinct clusters of PCa. **(a)** Disease-free survival outcomes for the six PCa clusters identified from 498 PCa patients from TCGA dataset (logrank-test p<0.0001), and **(b)** clusters reproduced by training a multi-class classifier and applying to an independent dataset (215 patients) from Taylors et al. (2010) (p=0.00088). Clusters C5 and C3 showed consistently poor prognosis in both the datasets.

Six individual classifiers were trained to classify each cluster *C_i_* **\in** {*C*_1_,..,*C*_6_} from the remaining five clusters to select genes (features) **F** for the combined (multiclass) classifier (see Methods). The individual classifiers achieved an AUC 0.853 on average, and 14 genes were selected into the multinomial (multi-class) classifier (**Table S2**). Five-fold cross-validation of the multinomial classifier trained on 80% TCGA data and tested on the remaining 20% data gave a max accuracy of 0.71 and average accuracy of 0.63. The classifier was then retrained using 100% TCGA data and applied on the independent Taylors et al. dataset to predict cluster labels C1-C6 (C1: 20, C2: 18, C3: 18, C4:9, C5: 39, C6: 36; total classified patients: 140). The predicted clusters showed distinct disease-free survival (p=0.00088) (**Figure 1b**) with C5 showing the quickest to relapse (>50% relapse by 72 months). Overall, the survival patterns of the clusters matched that observed for TCGA with C5 and C3 being the two worse survival clusters.

### PCa clusters display distinct metabolic profiles

We next computed the extent of deregulation of metabolic pathways-I in tumour clusters C1-C6 relative to normal tissues, using Pathifier (see Methods). Overall, carbohydrate and lipid metabolism pathways showed the most deregulation (elevated ‘activity’) in tumour samples relative to normals (**Figure 2a**), indicating increased energy requirement satisfied through increased metabolism of carbohydrates and lipids in the tumours. The elevated deregulation of the other three pathways, in particular, amino acid metabolism and TCA cycle (Kreb’s cycle; involves metabolism of acetyl-CoA derived from carbohydrates, fats and proteins to release energy, and provides important precurses for amino acid synthesis) in the tumour clusters suggests elevated synthesis and catabolism of amino acids, proteins, and nucleic acids via these pathways that is required for tumour cell growth and proliferation [24,25]. Note that all plots reported here have an overall ANOVA p-value ≤0.05 for differences between the cluster means.

**Figure 2:**
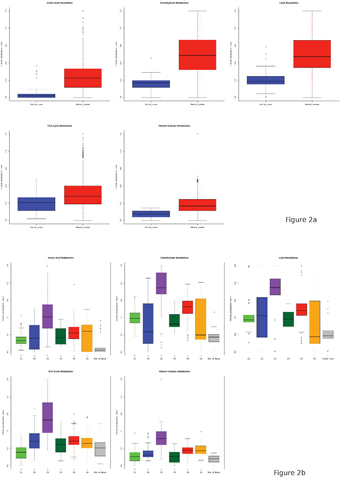
Deregulation of metabolic pathways-I. **(a)** The six patient clusters showed significantly elevated (p≤0.05) deregulation for all the five pathways from metabolic pathways-I relative to normal tissues, and in particular, for lipid and carbohydrate metabolism pathways. **(b)** The clusters displayed considerable heterogeneity among themselves, with the poor prognosis clusters C5 and C3 displaying the most deregulation for all the pathways.

While overall the tumour clusters showed elevated metabolism relative to normal, there was considerable heterogeneity among the tumour clusters, with the poor prognosis clusters C5, C3 and C6 showing the most deregulation for all pathways (**Figure 2b**). C2 and C6 showed the most ‘spread’ indicating heterogeneity even within these clusters. C3 stood out the most from the other clusters and showed a distinctly increased deregulation for the pathways, interestingly even for amino acid, TCA cycle and vitamin cofactor pathways, possibly suggesting a distinct metabolic profile for this cluster.

To further understand the high deregulation of carbohydrate and lipid metabolism pathways, we broke down the above analysis using metabolic pathways-II for the six tumour clusters (see Methods). Interestingly, among the carbohydrate metabolism pathways, we found the most deregulation for glycosaminoglycan synthesis and glycosaminoglycan degradation followed by pentose-phosphate metabolism (**Figure 3b**; larger versions available from Supplementary) whereas glycolysis (metabolism of glucose into pyruvate and energy) and galactose metabolism, which are thought to be more common forms of energy production [22,23], were relatively less deregulated in the tumour clusters (**Figure 3a**). Glycosaminoglycans are heteropolysaccharides and are among the key macromolecules that affect cell properties and functions, acting directly on cell receptors or via interactions with growth factors (see review [26]). Again, C3 and C5 and to an extent C2 and C6 showed the most deregulation for these pathways.

**Figure 3:**
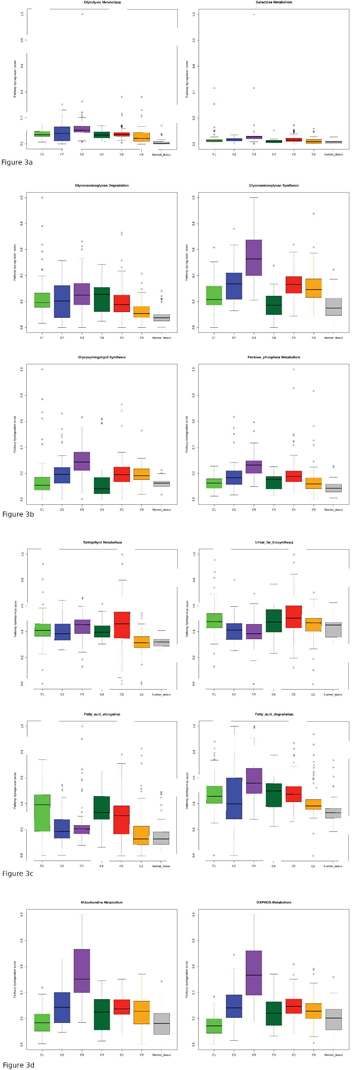
Deregulation of metabolic pathways-II. **(a)** Comparatively lower deregulation were observed for glucose (glycolysis) and galactose metabolism pathways. **(b)** On the other hand, al the clusters and particularly the poor-prognosis ones showed significantly elevated deregulation for sphingolipid and fatty-acid elongation and metabolism. **(c)** Similarly, the clusters also showed elevated deregulation for glycosaminoglycan, glycosphingolipid and pentose-phosphate metabolism. **(d)** Deregulation of these pathways was reflected as high activity of mitochondrial genes and the oxidative phosphorylation pathway.

Among the lipid metabolism pathways, the tumour clusters displayed elevated deregulation for sphingolipid and glycosphingolipid metabolism, with C5 and C3 showing the most deregulation for these pathways (**Figure 3c;** larger versions available from Supplementary). Sphingolipids are long-chain lipids and play important roles in signal transmission and cell recognition. Glycosphingolipids consist of sphingolipids with an attached carbohydrate. Sphingolipids and glycosphingolipids are believed to form mechanically stable and chemically resistant outer leaflet of the plasma membrane lipid bilayer [27] and have been thought to help cancer cells prevent drug intake through the membranes and thus become drug resistant [28]. Clusters C1, C4 and C5 showed elevated deregulation but C3 showed a relatively low deregulation for unsaturated fatty acid biosynthesis and fatty acid elongation; however, C3 (followed by C4 and C5) showed the most deregulation for fatty acid degradation, again highlighting the distinct metabolic expression profile for C3. Roles of fatty acid synthesis and elongation and fatty acid degradation in PCa has been noted in several studies (see reviews, [29] and [31]) which suggest increased expression of these mechanisms for membrane synthesis, storage and secretion, and to supply for the high energy needs in PCa relative to normal prostate tissues [30,31]. Finally, C3 and C5 showed the most deregulation for the pentose phosphatase pathway. This is a parallel metabolic pathway to glycolysis and it involves generation of pentoses and NADPH that is required for fatty acid synthesis [21].

We speculated that the increased deregulation of metabolism of the above reported complex forms of carbohydrates (glycosaminoglycans) and lipids (sphingolipids and glycosphingolipids), together with amino acid metabolism and TCA cycle would show up as increased deregulation of mitochondrial pathways. Accordingly, we found increased deregulation of mitochondrial subunits and the oxidative phosphorylation (OXPHOS) pathway (**Figure 3d**). OXPHOS constitutes a highly efficient mechanism of energy production in the mitochondria, and several recent studies indicate specific OXPHOS subtypes of cancers with elevated OXPHOS for energy production (shifting away or in addition to glycolysis) [35,36].

Collectively, the above results indicate that the tumour clusters: (i) are considerably heterogenous in their metabolic profiles (i.e., in their deregulation of metabolic pathways); (ii) depend on a combination of carbohydrate, lipid and protein/amino acid metabolism to meet their energy and cell-material synthesis needs; and (iii) based on the elevated deregulation for glycosaminoglycan, sphingolipid, glycosphingolipid metabolism and fatty acid degradation (compared to glycolysis), particularly for the poorer prognosis clusters C3, C5 and C6, the tumour clusters depend on synthesis and catabolism of more complex forms of carbohydrates and lipids through specific pathways in the mitochondria, and this can occur parallely or in the face of glycolysis for their energy production.

To understand the above differences in metabolic deregulation between the clusters in relation to cancer-cell survival, proliferation, and growth, we computed deregulation of two master regulator pathways that link cell metabolism to these processes namely, the AMPK/LKB1 and HIF1-a pathways [32,33,34]. The AMPK/LKB1 pathway includes LKB1, a master kinase that functions as a tumour suppressor and activates AMPK, a central metabolic sensor. AMPK regulates lipid and glucose metabolism in several metabolic tissues including the prostate [32]. The HIF1-a is a response pathway for hypoxia (low O_2_) and is a crucial survival pathway that enables adaptation to hypoxic conditions found especially in harsh tumour microenvironments ([33,34]). We observed significant deregulation of the two pathways across the different patient clusters (**Figure 4a&b**). For the AMPK/LKB1 pathway (**Figure 4a**), clusters C1 and C4 showed the most deregulation (higher activity), but interestingly C3, even though showed high deregulation but was in the opposite direction, that is, even lower activity than that of normal tissues. For the HIF-a pathway (**Figure 4b**), on the other hand, C3 and C5 showed the highest deregulation. We therefore speculate that the deregulation in metabolism for the poor prognosis clusters could be regulated or driven by lowered activity of the AMPK/LKB1 (in C3) and increased activity of the HIF-a pathway (in C3 and C5).

**Figure 4:**
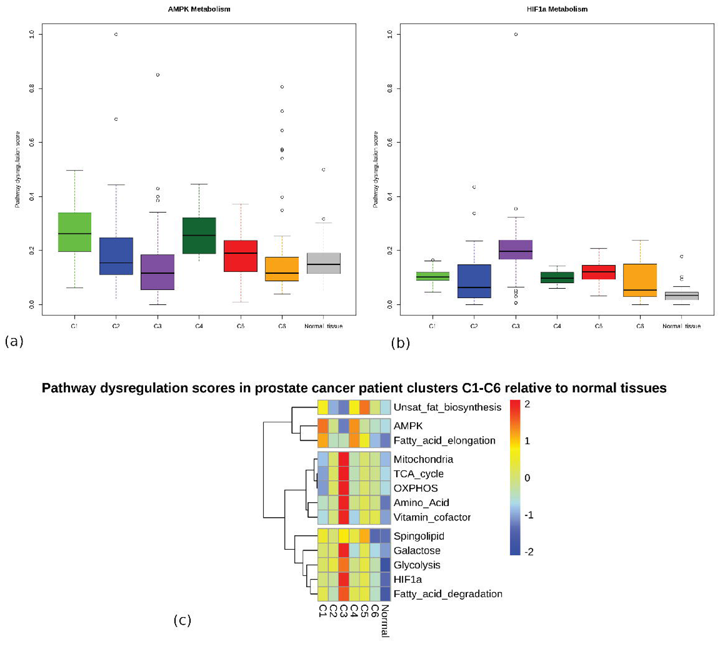
Deregulation of master regulators of metabolism. **(a)** AMPK/LKB1 pathway and **(b)** HIF1-a pathway. **(c)** A summary (showing average deregulation) of the pathways in the six clusters.

The above observations have been summarized in **Figure 4c**, where we show the mean deregulations for the above pathways for each patient cluster C1-C6.

### Enrichment for genetic alterations in the PCa clusters

Next, we sought to investigate whether any genetic changes in the form of DNA mutations, copy-number and mRNA-expression changes for known marker genes for PCa were enriched within the six PCa clusters. We assessed the status of 15 known predisposition genes or genes frequently altered in PCa namely, AR (androgen receptor), ATM, BRCA1, BRCA2, CDK12, MLH1, MSH2, MSH3, MSH6 (DNA double-strand break repair and/or mismatch repair), RB1 (cell cycle), TP53 (apoptosis response to DNA-damage and cell cycle regulation), SPOP, STK11 (LKB1, metabolism) and ERG-TMPRS22 (co-deletions). **Table 2** (and **Figure S2**) shows the enrichment for genetic alterations in these genes in the PCa clusters (see Methods). The table shows the clusters were enriched for alterations in different sets of genes, thus suggesting different genetic underpinnings for the clusters. For example, C3 was significantly enriched for alterations in STK11/LKB1 (p=1.31E-05), suggesting that the deregulation of the AMPK/LKB1 pathway in C3 that we observed previously (**Figure 4a**) could be driven by STK11/LKB1 alterations. Cluster C5, on the other hand, showed significant enrichment for defects in TP53 (p=4.37E-07), BRCA1 (p=8.6E-04), and MSH2 (p=3.2E-03), indicating that C5 could be underpinned by defects in the DNA-damage response (DDR). To confirm this observation, we computed the deregulation of DDR pathways using Pathifier (**Figure S3**), and indeed found the homologous recombination (HR) pathway to be substantially deregulated in C5 (C5 albeit also showed considerable heterogeneity for HR deregulation). Finally, cluster C1 showed significant (p=9.04E-05) enrichment for AR alterations (amplifications), indicating that C1 tumours could be primarily AR driven, whereas C4 appeared to be driven by ERG-TMPRSS2 deletions and fusions (p<0.04). In addition to these primary (highly significant) alterations, the clusters also showed moderate (but still significant, p≤0.05) enrichments for ATM (C1), PTEN (C1 and C5), RB1 (C3), SPOP (C3), and MSH2 (C5), thus suggesting that a combination of genetic alterations could be underpinning the phenotypes of the individual clusters. The enrichment for the above genetic alterations in patient clusters that were originally defined based solely on the expression of metabolic genes suggests that these genetic alterations, particularly in DDR genes, could be ‘pushing’ tumours towards adopting their specific ‘metabolic profiles’.

## Discussion

Here, we identified six clinically different (distinct disease-free survival) clusters of PCa patients based on clustering expression profiles of metabolic genes from TCGA prostate tissues (**Figure 1a**). The clusters were reproduced in an independent Taylors et al. tissue dataset (**Figure 1c**). We were next interested if any of the encoded proteins of these genes are also detectable from the blood and whether the blood expression profiles of those proteins are reflective of poor-prognosis PCa clusters. To test this possibility, we downloaded mRNA expression (microarray) data from the study by Olmos et al. (2012) [37] that measured expression profiles from whole blood samples from 42 good prognosis (selected for active surveillance; AS) and 65 advanced castrate resistant PCa (ACRPC) patients. We used our TCGA tissue-trained classifier (as before) to predict clusters (C1-C6) for Olmos et al. patients by plugging in the blood expression data of the corresponding proteins. **Table 3** shows the proportions (%) of AS and ACRPC patients labelled as C1-C6. Based on our TCGA tissue clustering (**Figure 1a**) if we consider clusters C1 and C4 as ‘good or better prognosis’ clusters and clusters C3 and C5 as ‘poor prognosis’ clusters, then we observe that 70.77% of ACPRC patients were categorized as ‘poor’ by the classifier, with most patients assigned to C3 (57%). However interestingly, 59.52% of AS patients were also (re)assigned to the poor clusters, with most to C3 again (35.71%). While these results are affected by differences between tissue and blood-based measurements – the blood circulates all throughout the body it is not representative of the prostate alone but is rather a ‘snapshot’ of the entire body – the results suggest that through a finer classification (six groups as against only AS vs ACPRC) based on metabolic profiling, we can better classify ‘grey-area patients’ and potentially reassign 35.71% AS patients to the poor-prognosis cluster C3. These results could mean that based on blood expression profiles, it may be hard to identify patients who ‘seem line’ AS but in fact show prognosis that of C3 – in other words, the C3 cluster of patients could have ‘distorted’ blood profiles because of which we may wrongly assign them to AS. While this is only our speculation, it is supported by the observation that even in the TCGA dataset, the time to biochemical relapse (BCR) as measured by rise in PSA levels measured from the blood does not correlate with actual time to relapse for the six clusters (**Figure 5**) – the poorest prognosis clusters C3 and C5 in fact show considerably late BCR compared to even the best prognosis clusters C1 and C4. We think that the ‘distorted’ PSA levels for C3 and C5 could be a result of deregulated metabolism, as recently been observed for obese men [6].

## Conclusions

The prostate is a metabolically active tissue. In this study, we identified six patient clusters of prostate cancer from TCGA and Taylors et al. datasets and these clusters showed distinct metabolic profiles and also distinct disease-free survival outcomes. We noted that the clusters were enriched for alterations in known PCa marker genes, indicating that that the alterations in these genes could be underpinning the PCa clusters and ‘pushing’ the clusters towards unique metabolic profiles. The metabolic link to PCa risk and aggressiveness is significant and has been investigated in several past [38,39] and recent [23, 31, 40] works, and some of these studies have identified obesity, high cholesterol, and visceral fat deposits around the prostate (peri-prostatic adipose tissue layer, PPAT layer) [42,43] as risk factors for PCa onset and aggressive progression. Towards this end, we are developing deep-learning models to analyse radiology (MRI and CT) scans from PCa patients to delineate the PPAT layer and build risk models for prediction of PCa aggressiveness. We foresee that accurate PPAT-layer delineation together with blood-based markers will lead us to a minimally invasive (radiology scans + blood test) approach for prediction of aggressive PCa.

---

Table 1: Clinical parameters for the six metabolic clusters (C1-C6) identified by clustering TCGA PCa data (498 patients) using 124 most altered metabolic genes (see Methods). The six clusters showed significantly different disease-free survival (**Figure 1a**), with a markedly poor-prognosis cluster C5 displaying relapse for >75% patients by 80 months. This cluster showed a markedly high PSA and high Gleason score.

Table 2: Enrichment (p-values) for genetic alterations in selected genes among patients in the six (C1-C6) TCGA clusters. Significant (p≤ 0.05) enrichments are in bold. We note that different clusters are enriched for alterations in different sets of genes – e.g. C1 for AR (p=9.04E-05), C3 for STK11 (p=1.31E-05) and RB1 (p=1.9E-04), and C5 for TP53 (p=4.37E-07) and BRCA1 (p=8.67E-04).

Table 3: Proportions (%) of active surveillance (AS) and advanced castrate-resistant prostate cancer (ACRPC) patients profiled in the study by Olmos et al. (blood expression) predicted for being in good (C1, C4) and poor (C3, C5) clusters. 70.77% of ACPRC patients were categorized as poor by the classifier, and interestingly 59.52% AS patients were reclassified as poor, with 35.71% assigned to C3.

